# ASCL1-regulated DARPP-32 and t-DARPP stimulate small cell lung cancer growth and neuroendocrine tumor cell survival

**DOI:** 10.1101/703975

**Authors:** Sk. Kayum Alam, Li Wang, Yanan Ren, Christina E. Hernandez, Farhad Kosari, Anja C. Roden, Rendong Yang, Luke H. Hoeppner

## Abstract

Small cell lung cancer (SCLC) is the most aggressive form of lung cancer, and new molecular insights are necessary for prognostic and therapeutic advances. Here we demonstrate in orthotopic mouse models that dopamine and cAMP-regulated phosphoprotein, Mr 32000 (DARPP-32) and its N-terminally truncated splice variant t-DARPP promote SCLC growth through increased proliferation, Akt/Erk-mediated survival and anti-apoptotic signaling. DARPP-32 and t-DARPP proteins are overexpressed in SCLC patient-derived tumor tissue, but virtually undetectable in physiologically normal lung. RNA sequencing analysis reveals a subset of SCLC patients with high tumoral t-DARPP expression and upregulated Notch signaling genes, including achaete-scute homologue 1 (ASCL1). We show that DARPP-32 isoforms are transcriptionally activated by ASCL1 in human SCLC cells. Taken together, we demonstrate new regulatory mechanisms of SCLC oncogenesis that suggest DARPP-32 isoforms may represent a negative prognostic indicator for SCLC and serve as a potential target for the development of new therapies.

## Introduction

Lung cancer is the leading cause of cancer death among both men and women in developed countries worldwide^1^. Small cell lung cancer (SCLC) kills approximately 250,000 people worldwide annually and comprises 15% of the incidences of lung cancer^2^. Cigarette smoking is the most common cause of SCLC, and SCLC patients have a smoking history in 95% of cases^3^. SCLC treatment options have not substantially changed since the introduction of cisplatin and etoposide decades ago^3^. SCLC is the most aggressive form of lung cancer due to its rapid doubling time and early widespread metastasis^4^. Most cases of SCLC are tumors of neuroendocrine origin with frequent inactivation of *TP53* and *RB1* as well as disruption of several molecular pathways, including Notch signaling^2^. SCLC patients typically present with advanced disease, respond to initial systemic chemotherapy, and then treatment refractory progression usually occurs within one year due to acquired drug resistance. Consequently, the median survival time of SCLC patients is only 9 to 20 months and merely 7% of SCLC patients survive beyond five years^4,5^. The frequent, rapid, and pronounced biological transition from chemotherapy-sensitive to chemotherapy-resistant SCLC underscores the importance of identifying therapeutically targetable molecular drivers of acquired resistance.

Dopamine and cyclic adenosine monophosphate-regulated phosphoprotein, Mr 32000 (DARPP-32) is an effector molecule that plays an important role in dopaminergic neurotransmission. Upstream of DARPP-32, dopamine D2 receptor agonists have been shown to inhibit lung tumor angiogenesis^6^, and clinical trials of selective dopamine D2 and D3 receptor antagonists have demonstrated anti-cancer efficacy in several cancer types other than lung^7^. Recent reports suggest aberrant DARPP-32 overexpression promotes oncogenesis in lung^8^, gastric^9^, colon^10^, prostate^11^, esophagus^12^, and breast adenocarcinomas^13^ through regulation of proliferation^14^, survival^15^, migration^8^, invasion^16^, and angiogenesis^17^. However, the role of DARPP-32 in neuroendocrine tumors remains unexplored. In the early 2000s, Wael El-Rifai and colleagues discovered that DARPP-32 and its novel transcriptional splice variant are frequently amplified and upregulated in gastric cancer^9,18^. The N-terminally truncated isoform of DARPP-32, named t-DARPP, uses a unique alternative first exon located within intron 1 of DARPP-32. DARPP-32 and t-DARPP are translated from a gene termed *phosphoprotein phosphatase-1 regulatory subunit 1B (PPP1R1B)* because full-length DARPP-32 inhibits protein phosphatase 1 (PP-1) activity following PKA-mediated phosphorylation at threonine-34 (T34) position. In turn, DARPP-32 inhibits PKA upon phosphorylation of its T75 residue by cyclin-dependent kinase 5 (Cdk5)^19^. Because t-DARPP lacks the first 36 amino acids of DARPP-32, including the T34 phosphorylation residue, t-DARPP is unable to inhibit PP-1^9^. Overexpression of t-DARPP in breast cancer has been shown to activate oncogenic PI3K/Akt signaling^20^. The dual function of DARPP-32 as either a kinase or a phosphatase inhibitor enables it to precisely modulate dopaminergic neurotransmission^19,21^ as well as regulate oncogenic signaling when its isoforms are aberrantly overexpressed in tumor cells.

We recently demonstrated that DARPP-32 and t-DARPP promote non-small cell lung cancer (NSCLC) growth in orthotopic mouse models, reduce apoptosis, activate Akt and Erk signaling, and enhance IKKα-mediated lung tumor cell migration^8^. Immunostaining of 62 human lung adenocarcinoma tissues showed that t-DARPP expression is elevated with increasing tumor staging score, a metric of tumor progression and growth. Bioinformatics analysis revealed upregulation of t-DARPP correlates with advanced tumor stage and poor overall survival of NSCLC patients^8^. Other groups have reported that t-DARPP promotes cancer cell survival by upregulation of Bcl2 in an Akt-dependent manner and causes drug resistance by activation of the Akt signaling pathway in breast cancer cells^15,22^. Studies have demonstrated that activation of Akt signaling by DARPP-32 and t-DARPP in breast and esophageal adenocarcinoma causes resistance to Herceptin (trastuzumab)^20,22–24^, a monoclonal antibody against HER2 commonly used in combination with chemotherapy to treat HER2-positive cancer. In breast cancer cells, DARPP-32 isoforms have been shown to promote resistance to lapatinib, a small molecule dual inhibitor of HER2/EGFR^13^, as well as EGFR inhibitors, erlotinib and gefitinib^25^. Most recently, it has been reported that activation of insulin-like growth factor-1 receptor (IGF1R) signaling in breast cancer cells initiates trastuzumab resistance by t-DARPP^26^. DARPP-32-mediated activation of IGF1R was also found to promote STAT3 signaling that contributes to gastric tumorigenesis^27^. Another study demonstrates that DARPP-32 interacts with ERBB3, activates Akt signaling, and exhibits resistance to gefitinib in esophageal adenocarcinoma^12^.

Given the multifaceted role of DARPP-32 and t-DARPP proteins in the oncogenesis of numerous cancer types^28^, including NSCLC^8^, we sought to determine whether DARPP-32 isoforms promote SCLC growth. Here, we demonstrate for the first time that DARPP-32 isoforms are regulated by Notch signaling to drive SCLC oncogenesis. Specifically, we show Notch target, achaete-scute homologue 1 (ASCL1), transcriptionally activates the DARPP-32 promoter. The pathogenesis of SCLC commonly involves inactivating Notch mutations, which lead to downstream ASCL1 activation^29^. Comprehensive genomic profiling of 110 SCLC patients revealed inactivating mutations in Notch family genes in 25% of human SCLC^30^. Correspondingly, activation of Notch signaling in a SCLC mouse model reduced tumor growth and extended survival of the mutant mice^30^. Concurrent Notch inactivation and ASCL1 overexpression in SCLC^31^ activates pulmonary neuroendocrine differentiation and stimulates tumor progression^32,33^. Our results support a model in which Notch inactivation leads to ASLC1-mediated activation of DARPP-32 isoforms, which we demonstrate promote tumor growth in an orthotopic mouse model of human SCLC by stimulating proliferation, Akt/Erk-mediated survival and anti-apoptotic signaling.

## Results

### DARPP-32 and t-DARPP promote small cell lung cancer growth

Numerous studies have reported the oncogenic role of DARPP-32 isoforms in breast and gastric malignancies^28,34^. Recently, we demonstrated overexpression of DARPP-32 isoforms in NSCLC promote tumor growth^8^. Based on these findings, we sought to investigate whether DARPP-32 isoforms contribute to the oncogenesis of SCLC. To begin to assess the role of DARPP-32 isoforms in SCLC, we transduced DMS-53 human SCLC cells with retroviral plasmids to overexpress exogenous DARPP-32, t-DARPP, or corresponding control LacZ protein (Fig. 1a). To determine how DARPP-32 isoform overexpression affects DMS-53 growth in vitro, we assessed cell confluency over time using the IncuCyte^®^ S3 live cell analysis system and found DARPP-32 isoform overexpression increases the growth rate of DMS-53 cells (Fig. 1b). We next sought to determine the effect of DARPP-32 isoforms on cell survival. Colorimetry-based MTS cell viability assays were performed using DMS-53 cells transduced with retrovirus overexpressing control or DARPP-32 isoforms. DMS-53 cells overexpressing DARPP-32 or t-DARPP protein exhibited increased viability compared to corresponding LacZ-transduced control cells (Fig. 1c). We replicated these studies using another human SCLC cell line, H1048. Stable retroviral overexpression of DARPP-32 and t-DARPP in H1048 cells (Fig. 1d) resulted in increased cell growth (Fig. 1e) and viability (Fig. 1f). Next, we silenced endogenous DARPP-32 protein expression through lentiviral shRNA-mediated transduction in DMS-53 cells using two different shRNAs targeting distinct regions of DARPP-32 (Fig. 1g). In accordance with our DARPP-32 overexpression findings (Fig. 1a-f), we observed decreased DMS-53 growth (Fig. 1h) and reduced cell viability (Fig. 1i) upon DARPP-32 protein ablation using live cell analysis and viability assays, respectively. Similarly, shRNA-mediated DARPP-32 protein knockdown in H1048 cells (Fig. 1j) resulted in a reduced cellular growth rate (Fig. 1k) and fewer viable cells (Fig. 1l) relative to corresponding LacZ shRNA transduced control SCLC cells. Based on our cumulative findings, we conclude that DARPP-32 and t-DARPP positively regulate cell growth and survival in SCLC cells.

**Fig. 1:**
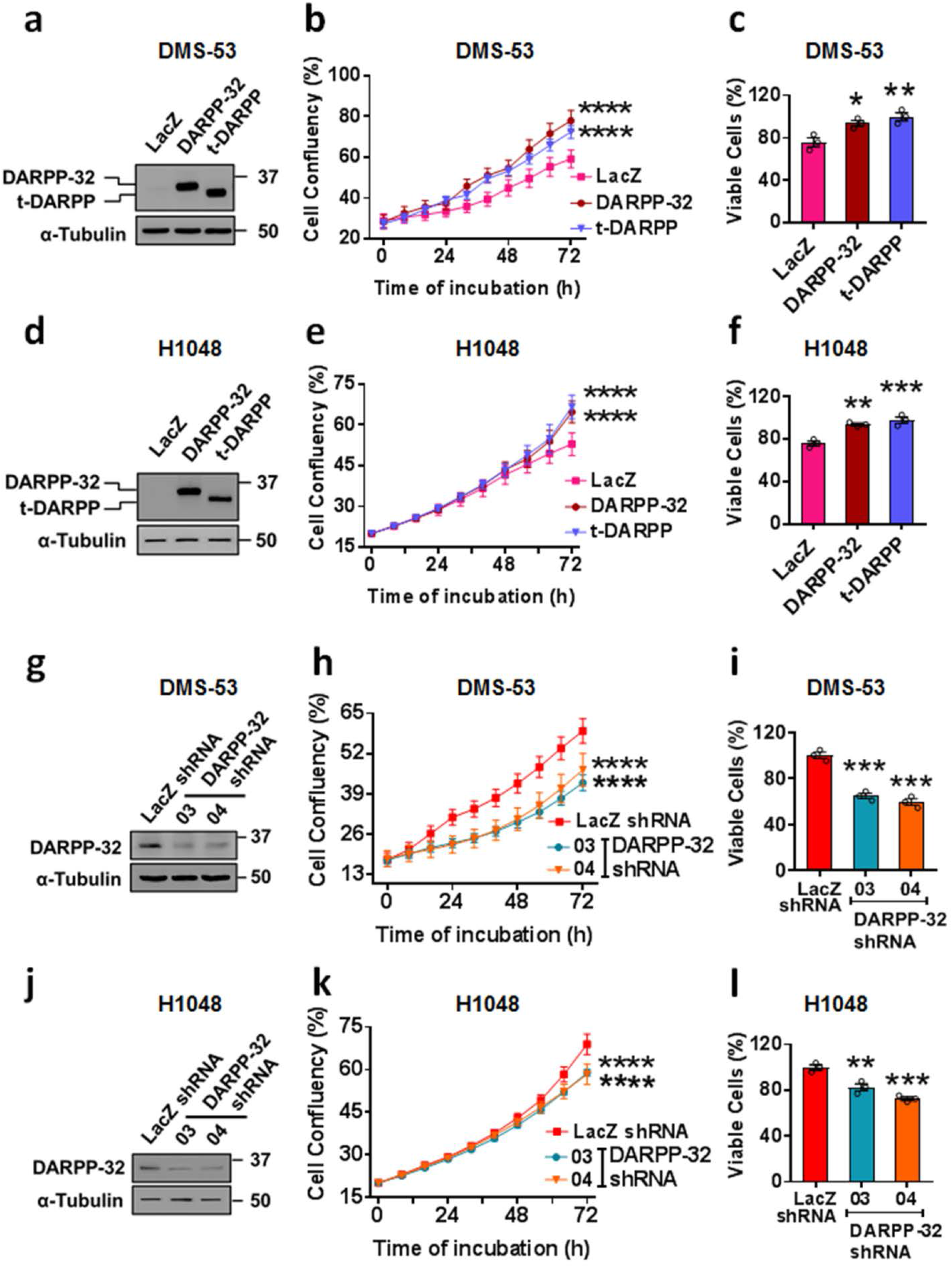
DARPP-32 isoforms promote cell growth and survival. **a** Retrovirus encoding control- (LacZ), DARPP- 32- or t-DARPP-overexpressing clones were transduced in human DMS-53 SCLC cells. Immunoblotting of cell lysates was performed to detect DARPP-32, t-DARPP and α-tubulin (loading control) proteins. **b** DMS-53 cells expressing control-, DARPP-32- or t-DARPP-overexpressing clones were seeded into 96-well cell culture plates. Images were captured at every 6h interval using IncuCyte^®^ live cell analysis imaging system. Cell confluency was determined by measuring the area occupied by the cells in each well. **c** DMS-53 cells transduced with control-, DARPP-32- or t-DARPP-overexpressing clones were plated into 96-well cell culture plates for 72h. MTS-1 reagents were used to quantify viable cells by measuring absorbance at 490 nm. **d** DMS-53 cells were transduced with lentivirus encoding control shRNA (LacZ) or DARPP-32 shRNAs (clone numbers 03 and 04). Immunoblot assays were performed to detect DARPP-32 and α-tubulin proteins. **e** DMS-53 cells transduced with control shRNA or DARPP-32 shRNAs were seeded into 96-well cell culture plates. IncuCyte^®^ live cell analysis imaging system was used to determine cell confluency by measuring the cell-occupied area. **f** Colorimeter-based cell survival assays using MTS reagents were performed in DMS-53 cells transduced with control shRNA or DARPP-32 shRNAs. **g** Human H1048 SCLC cells transduced with control-, DARPP-32- or t-DARPP-overexpressing clones were lysed and subjected to immunoblot using DARPP-32 and α-tubulin (loading control) antibodies. **h** H1048 cells were transduced with retrovirus encoding control-, DARPP-32- or t-DARPP-overexpressing clones. The cell-occupied area in each well was assessed using the IncuCyte^®^ live cell analysis imaging system. **i** Cell survival assay of H1048 cells transduced with retrovirus containing control- (LacZ), DARPP-32- or t-DARPP-overexpressing clones was performed using MTS-1 reagent. **j** H1048 cells transduced with lentivirus encoding control or DARPP-32 shRNAs were plated in 60mm cell culture dishes. DARPP-32 and α-tubulin (loading control) proteins were detected by immunoblotting of cell lysates. **k** Cell confluency of H1048 cells transduced with control or DARPP-32 shRNAs was measured using IncuCyte^®^ live cell analysis imaging system. **l** H1048 cells were transduced with lentivirus encoding control or DARPP-32 shRNAs. Colorimeter-based in vitro assay was performed to measure cell survival using MTS-1 reagents. Each open circle on a bar graph represents an independent experiment. Immunoblots are representative of three independent experiments. Error bars indicate SEM (n=3). \**P*<0.05, \*\**P*<0.01, \*\*\**P*<0.001 and \*\*\*\**P*<0.0001, one-way ANOVA followed by Dunnett’s test for multiple comparison.

### DARPP-32 isoforms negatively regulate apoptosis

Given that DARPP-32 isoforms promote SCLC cell growth and survival, we sought to determine the role of DARPP-32 isoforms regulating apoptosis in SCLC cells. We first performed flow cytometry-based annexin V apoptosis assays in DMS-53 cells retrovirally transduced with control-, DARPP-32- or t-DARPP- overexpressing vectors. We observed a reduced number of annexin V-positive cells (Fig. 2a: lower right quadrant; annexin V-FITC positive and propidium iodide negative) in DARPP-32- and t-DARPP-overexpressing DMS-53 cells, which suggests DARPP-32 isoform overexpression causes a reduction in apoptosis (Fig. 2a: right). We next assessed apoptosis in DARPP-32-depleted DMS-53 cells using flow cytometry-based annexin V assays. As expected, DARPP-32 ablation caused an increase in annexin V-positive cells relative to controls (Fig. 2b). To determine how DARPP-32 regulates the expression of apoptosis-associated proteins, we performed immunoblotting studies in human DMS-53 and H1048 SCLC cells overexpressing DARPP-32 isoforms or corresponding control LacZ. Briefly, caspase proteins are cysteine-dependent aspartate-specific proteases that play central role in mediating apoptosis. An initiator caspase (e.g. Caspase-8, −9) proteolytically cleaves an effector caspase (e.g. Caspase-3, −6) upon binding to specific oligomeric activator protein (e.g. Apaf-1). The active effector caspases then proteolytically degrade a host of intracellular proteins (e.g. PARP-1) to carry out the cell death program^35^. We observed reduced expression of cleaved poly(ADP-ribose) polymerase-I (PARP-I) and caspase-3 proteins in DARPP-32- and t-DARPP-overexpressing cell lines relative to controls (Fig. 2c and d), suggesting DARPP-32 isoforms promote apoptosis. Conversely, shRNA-mediated depletion of DARPP-32 isoforms resulted in decreased caspase-3 and PARP-I cleavage in DMS-53 (Fig. 2e) and H1048 (Fig. 2f) human SCLC cells. Collectively, our findings suggest DARPP-32 upregulation correlates with increased apoptosis in human SCLC cells.

**Fig. 2:**
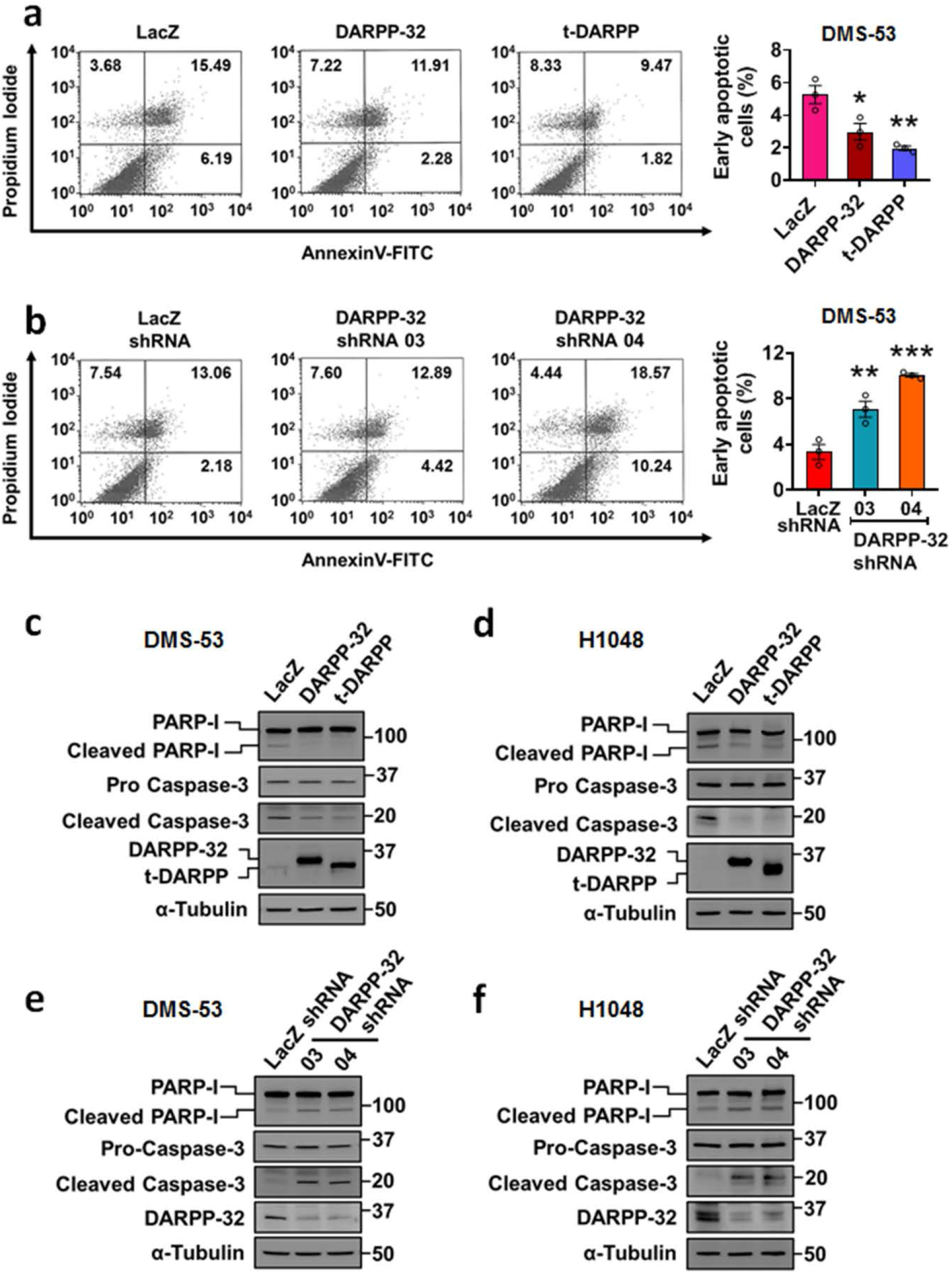
DARPP-32 and t-DARPP decreases cell death by promoting apoptosis. **a** Retrovirus containing control- (LacZ), DARPP-32- or t-DARPP-overexpressing clones and **b** lentivirus encoding control (LacZ) or DARPP-32 shRNAs were transduced in DMS-53 cells. Cells were incubated with anti-annexin V antibodies conjugated with FITC followed by propidium iodide incorporation. Flow cytometry-based apoptosis assays were performed to determine the total number of annexin V-positive cells. The average number of annexin V-positive cells of three independent experiments were plotted in a bar graph. Each open circle on a graph represents an independent experiment. The numerical values on quadrants of the scatter plots represent the percentage of total cells in one single representative experiment. **c** DMS-53 and **d** H1048 cells were transduced with control-, DARPP-32- or t-DARPP-overexpressing clones. Cell lysates were collected and immunoblotted with antibodies to detect cleaved and uncleaved PARP-I, cleaved and uncleaved (i.e., pro-) caspase-3, DARPP-32 and α-tubulin (loading control). **e** DMS-53 and **f** H1048 cells were transduced with lentivirus encoding control or DARPP-32 shRNAs. Cleaved and uncleaved PARP-I, cleaved and uncleaved (i.e., pro-) caspase-3, DARPP-32 and α-tubulin (loading control) proteins were detected by immunoblotting of cell lysates. All bar graphs represent mean ± SEM (n=3). \**P*<0.05, \*\**P*<0.01 and \*\*\**P*<0.001, one-way ANOVA followed by Dunnett’s test for multiple comparison.

### Akt/Erk-mediated cell proliferation is regulated by DARPP-32 in SCLC

Based on prior findings that Akt and Erk1/2 signaling pathways contribute to DARPP-32 mediated breast and gastric oncogenesis, we sought to elucidate their potential role in SCLC tumor growth. We first evaluated the phosphorylation status of Akt and Erk in human SCLC cells by immunoblotting. Stable retroviral overexpression of exogenous DARPP-32 and t-DARPP substantially enhanced phosphorylation levels of Akt and Erk in DMS-53 and H1048 cells (Fig. 3a-b). Corresponding total Akt and Erk1/2 protein levels in SCLC cells overexpressing DARPP-32 isoforms remained consistent by immunoblotting (Fig. 3a-b). We next performed immunoblotting studies using human DMS-53 and H1048 SCLC cells transduced with lentivirus encoding DARPP-32 shRNAs or control LacZ shRNA. We found ablation of endogenous DARPP-32 decreases phosphorylation levels of Akt and Erk1/2 relative to LacZ shRNA-transduced controls, while the expression of corresponding total Akt and Erk1/2 protein remains unchanged upon DARPP-32 modulation (Fig. 3c-d). Given Akt and Erk activation promote cell proliferation^8^, we next performed flow cytometry-based BrdU cell proliferation assays in DMS-53 cells transduced with DARPP-32 or control shRNA. We observed reduced BrdU staining in DARPP-32-depleted DMS-53 cells (Fig. 3e), suggesting DARPP-32 isoforms directly regulate human SCLC cell proliferation via Akt/Erk signaling.

**Fig. 3:**
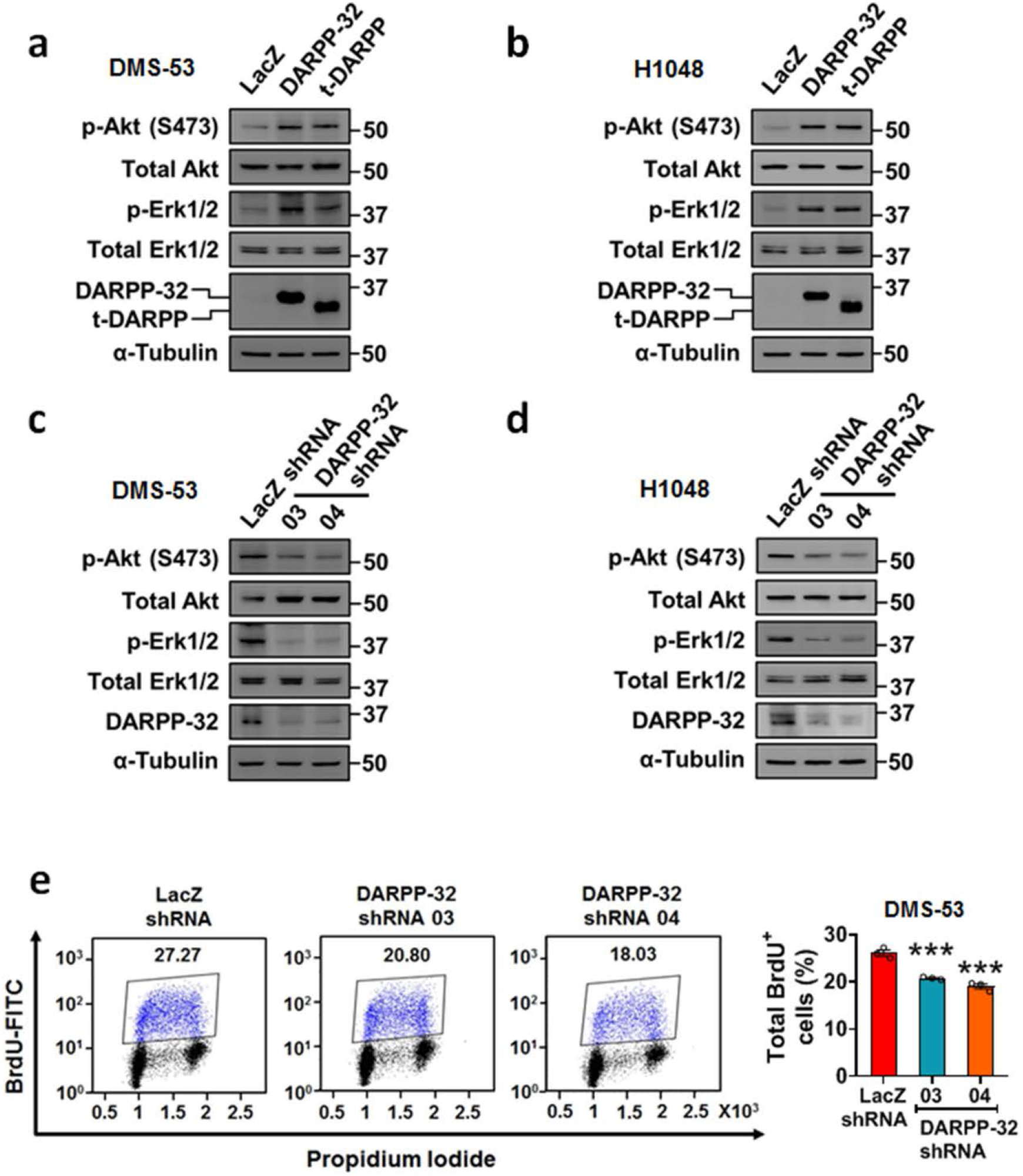
DARPP-32 and t-DARPP positively regulates cell survival through Akt and Erk1/2. **c** Lysates of DMS-53 and **d** H1048 cells overexpressing control, DARPP-32 or t-DARPP clones were immunoblotted using antibodies against phosphorylated Akt (p-Akt; S473), total Akt, phosphorylated Erk1/2 (p-Erk1/2, T202/Y204), total Erk1/2, DARPP-32 and α-tubulin (loading control). **e** DMS-53 and **f** H1048 cells transduced with control or DARPP-32 shRNAs were subjected to western blotting using antibodies against phosphorylated Akt (p-Akt; S473), total Akt, phosphorylated Erk1/2 (p-Erk1/2), total Erk1/2, DARPP-32 and α-tubulin (loading control).

### DARPP-32 and t-DARPP promote SCLC growth in orthotopic mouse xenograft models

Based on our results that SCLC cell growth, survival, and proliferation is regulated by DARPP-32 isoforms, we sought to determine whether DARPP-32 drives SCLC growth in vivo. To this end, we utilized an orthotopic lung cancer xenograft mouse model^8^. Specifically, we injected luciferase-labeled DARPP-32-depleted human DMS-53 SCLC cells into the left thorax of anesthetized SCID mice. After establishment of the lung tumor, we xenogen imaged the mice regularly over the course of five weeks. Mice challenged with DARPP-32-ablated DMS-53 cells showed a substantial decrease in lung tumor growth compared to mice challenged with human SCLC cells transduced with control LacZ shRNA (Fig. 4a) suggesting that DARPP-32 promotes human SCLC growth in mouse xenograft models. We next sought to determine whether overexpression of DARPP-32 isoforms promotes SCLC tumor growth in vivo. Briefly, 1×10^6^ luciferase-labeled human DMS-53 SCLC cells stably overexpressing exogenous DARPP-32 or t-DARPP were injected into the left thorax of anesthetized SCID mice, and we observed increased tumor growth in mice harboring DMS-53 cells overexpressing DARPP-32 or t-DARPP relative to mice that received control DMS-53 cells overexpressing LacZ (Fig. 4b). Correspondingly, we demonstrated increased tumor growth in mice challenged with H1048 cells overexpressing DARPP-32 and t-DARPP proteins (Fig. 4c). Taken together, our data suggest DARPP-32 isoforms promote SCLC growth in orthotopic mouse xenograft models.

**Fig. 4:**
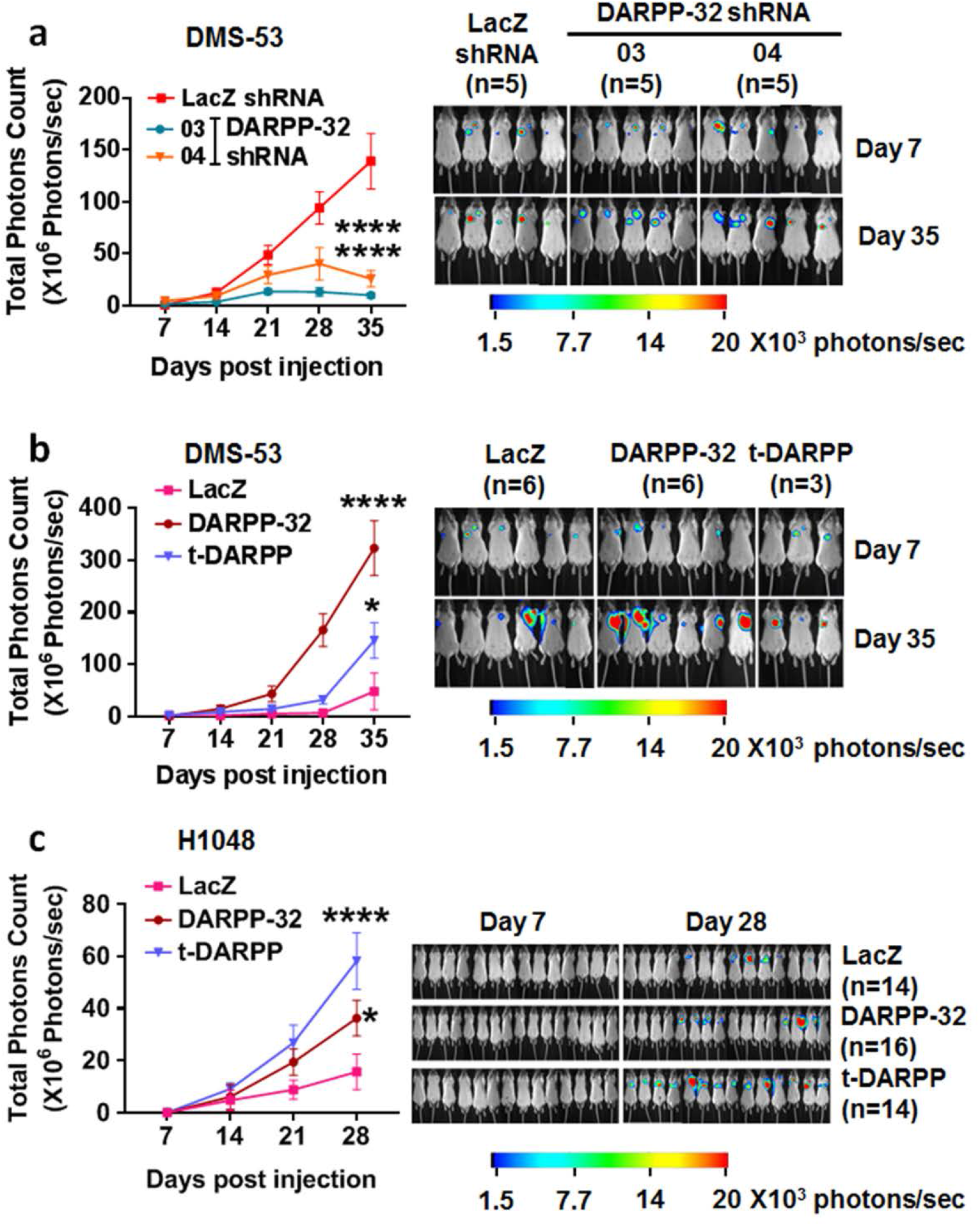
DARPP-32 isoforms promote growth of human SCLC cells in mouse xenograft models. **a** Luciferase-labeled human DMS-53 cells transduced with lentivirus encoding control (LacZ) or DARPP-32 shRNAs were orthotopically injected into the left thorax of SCID mice and imaged for luminescence on indicated days. **b** Luciferase-labeled human DMS-53 and **c** H1048 cells overexpressing control (LacZ), DARPP-32 or t-DARPP clones were injected into the left thorax od SCID mice and imaged for luminescence on the indicated days. In vivo images of SCID mice bearing luciferase-labeled tumors showing luminescence on the indicated days are depicted. Total luminescence intensity (photons count) was calculated using molecular imaging software. The numerical value of luminescence was represented by the color bar. Error bars indicate SEM. \**P*<0.05 and \*\*\*\**P*<0.0001, one-way ANOVA followed by Dunnett’s test for multiple comparison

### Elevated DARPP-32 expression is linked to small cell lung tumorigenesis

Given that DARPP-32 ablation reduces tumor growth in mouse models of human SCLC, we aimed to elucidate the clinical relevance of DARPP-32 in SCLC patients. Based on published reports that indicate upregulation of DARPP-32 and t-DARPP have been associated with breast, gastric, colorectal and non-small cell lung cancer^28,34^, we sought to assess DARPP-32 and t-DARPP protein expression in SCLC patients. Immunohistochemistry was performed to detect DARPP-32 isoforms in tissue specimens obtained from small cell lung carcinoma patients. Briefly, we obtained serial whole tissue sections of formalin-fixed paraffin embedded tumor tissue blocks corresponding to each patient and immunostained with an antibody that detects both DARPP-32 and t-DARPP via a C-terminal epitope present in both isoforms. Using the same antibody, we also immunostained normal human lung tissue specimens. We observed strong expression of DARPP-32 isoforms in human SCLC specimens, whereas DARPP-32 and t-DARPP protein expression was virtually undetectable in physiologically normal human lung tissue (Fig. 5). The percentage of tumor cells positive for DARPP-32 isoforms and their staining intensity was scored by a pulmonary pathologist (ACR) using a scale of 0-3 (i.e. 0= none, 1= weak, 2= moderate, 3= strong expression). Based on the percentage of tumor cells staining positive and the staining intensity in those cells, we calculated an IHC score (IHC score = percentage of tumor cells x staining intensity) for each specimen using the pathological scoring (Supplementary Table 1). In summary, our immunostaining studies suggest DARPP-32 isoforms are aberrantly overexpressed in human SCLC.

**Fig. 5:**
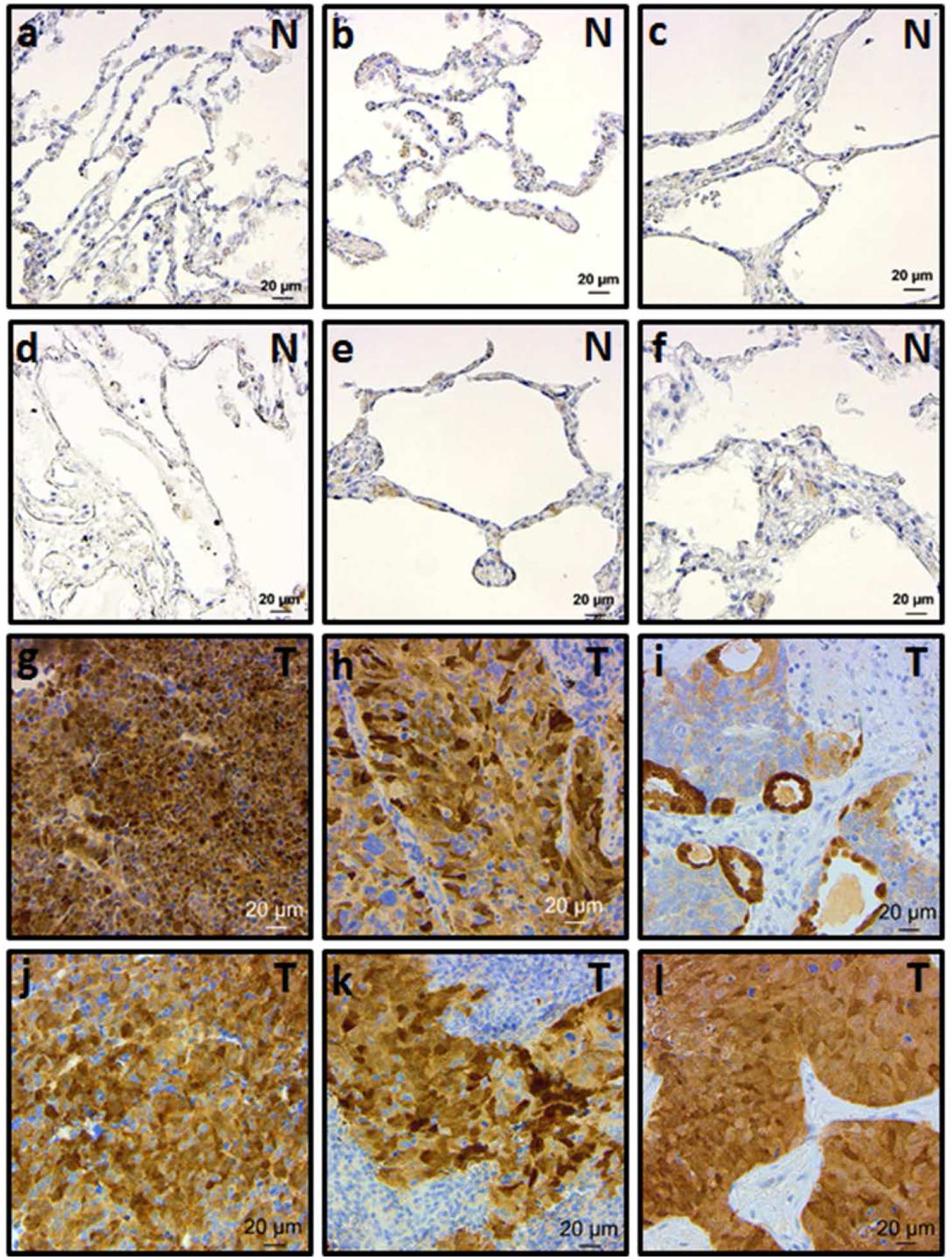
Expression of DARPP-32 isoforms elevated in human SCLC patients. **a-f** Normal lung tissues and **g-l** tumor tissues from SCLC patients were subjected to immunohistochemistry using antibodies that recognize both DARPP-32 and t-DARPP isoforms. Nuclei were counterstained using hematoxylin stain. N and T denotes normal and tumor, respectively. Scale bar indicates 20μm.

### Upregulated t-DARPP protein regulates Notch signaling

We sought to elucidate the molecular mechanisms through which DARPP-32 isoforms promote human SCLC. First, we assessed DARPP-32 and t-DARPP transcript levels in a RNA-Seq dataset consisting of 29 human SCLC patients, 23 of which also contained corresponding matched normal lung tissue^36^. We found that t-DARPP transcripts were elevated in SCLC relative to normal lung tissue (Fig. 6a). Interestingly, we identified six SCLC patients within this cohort with high levels of t-DARPP transcript in tumor tissue, but undetectable t-DARPP in corresponding normal tissue (blue data points in Fig. 6a). Using differential gene expression to analyze the high tumoral t-DARPP subset, we identified 3698 upregulated genes and 3004 downregulated genes in tumor relative to matched normal tissue (Fig. 6b and Supplementary Table 2-3). We employed gene set enrichment analysis (GSEA) and identified five cellular pathways regulated by t-DARPP in human SCLC (Fig. 6c, Supplementary Figure 1, Supplementary Table 4-5). We chose to focus our investigation on how Notch signaling is regulated by t-DARPP based on the well-established role of Notch family members in SCLC growth. To that end, we examined differential tumor vs. normal expression of the 31 Notch family members that comprised the gene set (Supplementary Table 4). As expected, achaete-scute homologue 1 (ASCL1), a main driver of SCLC, was increased nearly 50-fold in tumor tissues compared to matched normal specimens (Fig. 6d). Physiologically, ASCL1 is predominantly expressed in neural progenitor cells and supports growth and development of nervous tissue^37^. Pathologically, ASCL1 is overexpressed in aggressive neuroendocrine lung cancers and linked to SCLC progression^38^. The transcription factor ASCL1 activates Notch signaling by directly inducing expression of the ligands Delta and Jagged, thereby promoting oncogenesis in SCLC as well as T cell acute lymphoblastic leukemia (T-ALL)^39^. We sought to identify target genes controlled by DARPP-32 isoforms through transcriptional regulation by ASCL1 that might contribute to SCLC growth and progression. We identified 149 such target genes (Supplementary Table 6) common between 456 ASCL1 targets found through ChIP-seq analysis^40^ and 6702 genes (Supplementary Table 2-3) that were differentially regulated in SCLC patients^36^ with high tumoral t-DARPP expression relative to matched normal samples (Fig. 6e). We selected 16 genes that had been implicated in cancer and ASCL1-regulated Notch signaling for further confirmatory analysis. Using qRT-PCR, we compared transcript levels of each gene in DARPP-32 shRNA-versus LacZ control shRNA-transduced DMS-53 human SCLC cells. We found that mRNA expression of 10 genes, including ASCL1, was substantially reduced in DARPP-32-depleted cells compared to LacZ shRNA-transduced controls (Fig. 6f).

**Fig. 6:**
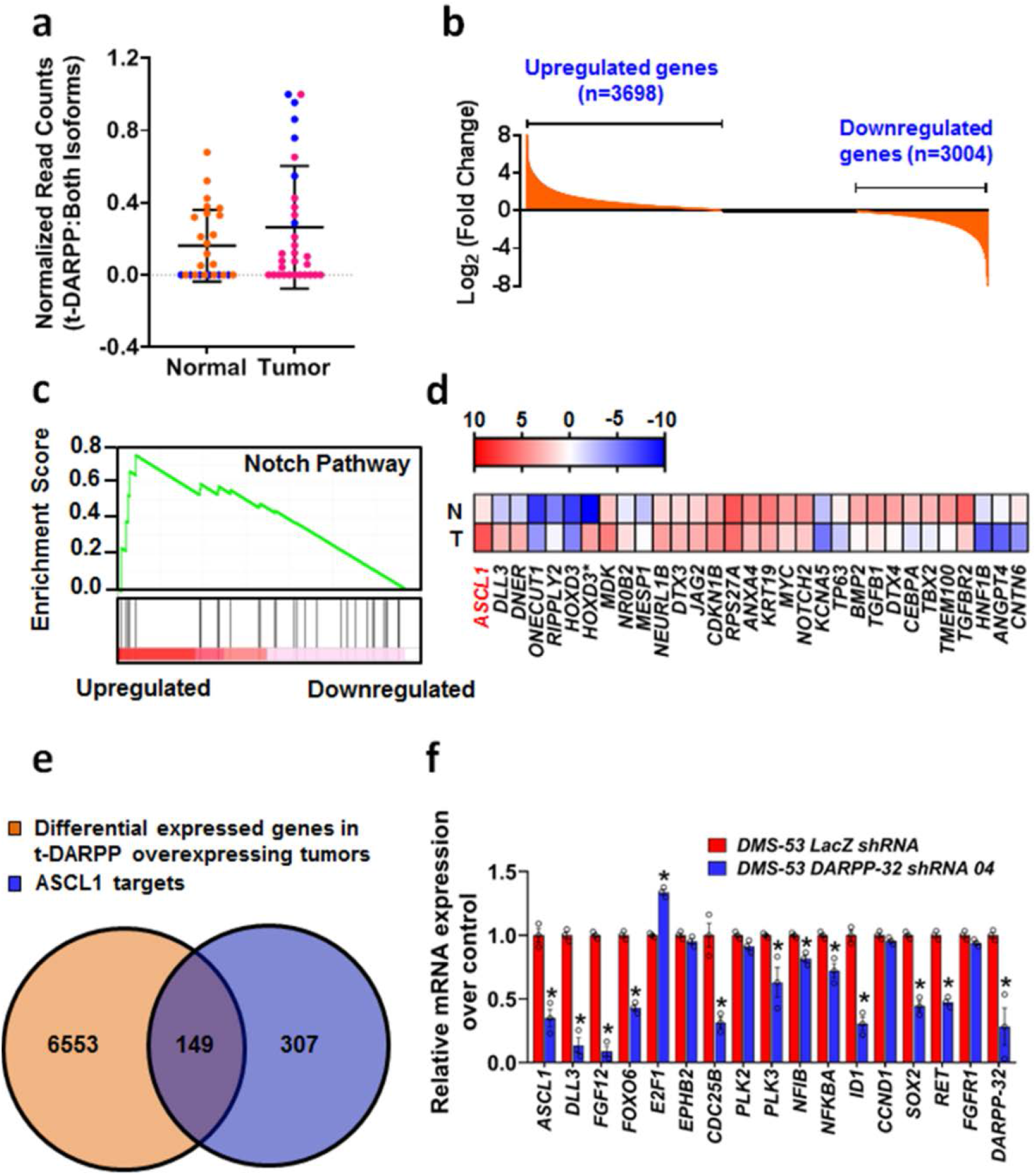
Elevated t-DARPP regulates Notch signaling through ASCL1 in SCLC. **a** Expression of t-DARPP transcripts was quantified using previously published RNA-Seq database^36^. Blue points indicate individual SCLC patients (n=6) with an elevated t-DARPP expression (t-DARPP: both DARPP-32 isoforms) in tumor tissue compared to mRNA derived from corresponding adjacent normal tissue. **b** Quantification of mRNA transcripts in a subset of SCLC patients (n=6; blue points) expressing high level of t-DARPP isoforms in tumor relative to normal tissue. **c** Gene set enrichment analysis (GSEA) was performed to identify cellular pathways regulated in a subset of SCLC patients with elevated t-DARPP levels in tumor tissue. Notch signaling was enriched by GSEA. **d** Heat map showing the expression of Notch family member genes identified through GSEA. N and T denotes normal and tumor tissue, respectively. Red color represents highest fold change in expression in tumor tissue relative to normal. The asterisk indicates an alternative splice variant of a gene. Color bar represents normalized read counts in a log scale. **e** Venn diagram exhibiting the number of differential expressed genes in a subset of patients expressing elevated t-DARPP transcripts and previously reported ASCL1 targets^40^. **f** Sixteen targets (selected from those represented in the union of the Venn diagram in e) were quantified using qRT-PCR in DMS-53 cells transduced with lentivirus encoding control (LacZ) or DARPP-32 shRNAs. Each open circle on a bar graph denotes three independent experiments and error bars represent SEM. \**P*<0.05, two-way ANOVA followed by Sidak's multiple comparisons test.

### ASCL1 transcriptionally regulates DARPP-32 expression

Given that most of SCLCs overexpress ASCL1 protein^41^, we sought to determine the role of ASCL1 in regulating DARPP-32 expression. We first performed immunoblotting experiments to assess endogenous expression of ASCL1 in DMS-53 and H1048 cells. Based on our results, we classified DMS-53 and H1048 cells as ASCL1-positve and −negative cells, respectively (Fig. 7a). To address whether ASCL1 regulates DARPP-32 promoter activity, we first stably depleted ASCL1 protein expression in DMS-53 cells via lentiviral-mediated shRNA transduction (Fig. 7b). We next determined DARPP-32 promoter activity by performing dual-luciferase assays in DMS-53 cells transduced with either ASCL1 shRNA or control (LacZ) shRNA. Briefly, DARPP-32 promoter (−3115 to −95 from transcription start site) was previously cloned into pGL3 luciferase reporter vector and characterized by El-Rifai et al^42^. We transfected DMS-53 cells with pGL3-DARPP-32 (firefly luciferase) and pRL-SV40 (renilla luciferase; acts as transfection control) plasmids to assess DARPP-32 promoter activity by measuring luminescence at 48h post-transfection. We observed a reduction in DARPP-32 promoter activity in ASCL1 ablated DMS-53 cells (Fig. 7c). To understand how ASCL1 regulates DARPP-32 expression, we individually mutated each of three putative ASCL1 binding sites on DARPP-32 promoter (i.e. CAGCTG motif at −1758 to −1752, −1648 to −1642, and −1263 to −1257) using site-directed mutagenesis approach. We also generated every combination of double and triple mutated ASCL1 binding sites to net a total of seven ASCL1 binding site mutants (Fig. 7d). To test whether the putative ASCL1 binding sites on the DARPP-32 promoter are required for ASLC1-mediated transcriptional activation of DARPP-32, we performed dual luciferase assays in DMS-53 cells transfected with either wild-type (pGL3-DARPP-32; clone A) or ASCL1 binding site-mutated (clones B to H) DARPP-32 promoter constructs (Fig. 7d). Mutation of any of the ASCL1 binding sites reduced DARPP-32 promoter activity (Fig. 7e), suggesting all three sites are involved and necessary for ASCL1 regulation of DARPP-32. To further confirm that ASCL1 transcriptionally regulates DARPP-32, we next performed dual luciferase assays in ASCL1-negative H1048 cells in which we exogenously overexpressed ASCL1 (Fig. 5f). As expected, ASCL1 overexpression in H1048 cells increases wildtype DARPP-32 promoter (A) activity, whereas exogenous ASCL1 has little effect on its mutant counterparts (B to H; Fig. 5g). Taken together, we identify three novel ASCL1 binding sites on the DARPP-32 promoter through which ASCL1 transcriptionally activates DARPP-32.

**Fig. 7:**
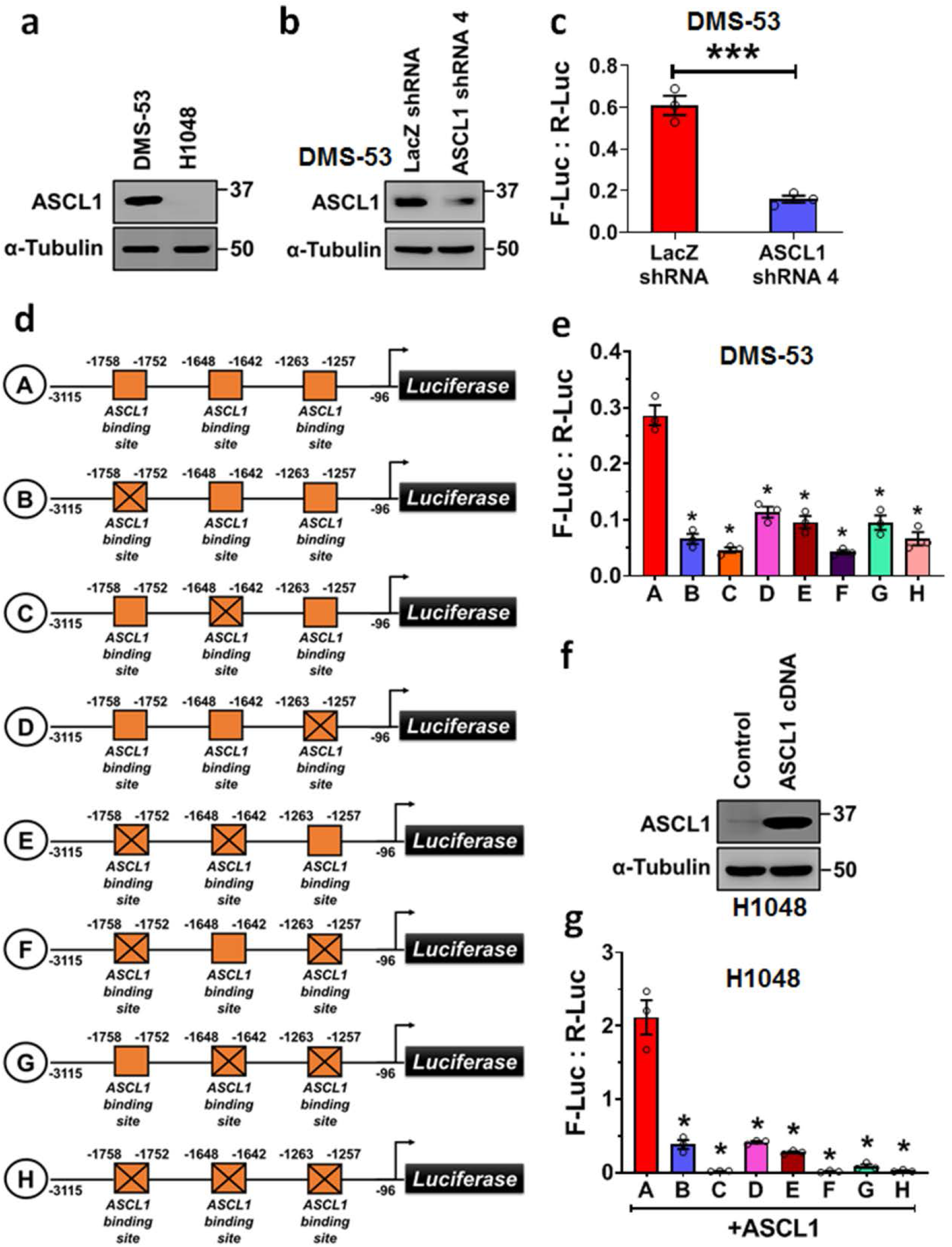
ASCL1 positively regulates DARPP-32 expression in SCLC cells. **a** Two SCLC cell lines were lysed and immunoblotted with antibodies against ASCL1 and α-tubulin (loading control). **b** DMS-53 cells transduced with control (LacZ) or ASCL1 shRNAs. Cell lysates were collected and subjected to western blot to measure the knockdown of ASCL1. **c** Dual-luciferase assays were performed in DMS-53 cells transduced with either control or ASCL1 shRNAs following transient transfection of pGL3-DARPP-32-Luc (Firefly) and pRL-SV40 (Renilla) plasmids. Firefly and renilla luminescence was measured and plotted as ratio. Mean ±SEM (n=3), two-tailed unpaired t-test. **d.** Three potential ASCL1 binding sites are present within the DARPP-32 promoter (yellow boxes). ASCL1 binding sites on the DARPP-32 promoter have been mutated by site-directed mutagenesis. The crossed boxes refer to the mutated sites. **e.** DMS-53 cells transfected with either site-directed mutagenesis clones (indicated as B to H) or pGL3-DARPP-32-Luc plasmids (A) were subjected to dual-luciferase assay. **f.** Immunoblot was performed in H1048 cells transfected with ASCL1 expressing plasmids (cDNA). **g.** Transient transfection of either site-directed mutagenesis clones (indicated as B to H) or pGL3-DARPP-32-Luc plasmids (A) was carried out in H1048 cells exogenously overexpressed ASCL1. Dual-luciferase assays were performed after 48h post-transfection. Each open circle on a graph represents an independent experiment. Results represent mean ±SEM (n=3). *P<0.0001, one-way ANOVA followed by Dunnett’s test for multiple comparison.

## Discussion

While growing evidence supports the oncogenic role of DARPP-32 and t-DARPP in a wide variety of adenocarcinomas^28,34^, we are the first to report that DARPP-32 isoforms promote neuroendocrine oncogenesis, as we observe DARPP-32 and t-DARPP drive SCLC growth through anti-apoptotic mechanisms and pro-survival signaling via Akt and Erk. Neuroendocrine tumors arise from cells of endocrine and neural origin and have been reported in the pancreas^43,44^, gastrointestinal tract^43,44^, lung^45,46^, breast^47^, genitourinary tract^48^, liver^49^, gall bladder^50^, and glands including thymus^51^. Pulmonary neuroendocrine tumors are categorized as low-grade and high-grade based on clinical and molecular factors^52^. SCLCs and large cell neuroendocrine carcinomas comprise high-grade neuroendocrine tumors of the lung^53^, which are characterized by a strong association with smoking, high mitotic indices, rapid growth, metastasis, frequent *TP53* and *RB1* mutations, and poor survival^2^. High grade neuroendocrine tumors frequently exhibit Notch inactivation and overexpression of achaete-scute homolog 1 (ASCL1), a transcription factor that functions as a master regulator of neuroendocrine programming and acts as a lineage-specific oncogene^29^. Here, we show that ASCL1 transcriptionally activates DARPP-32 isoforms in human SCLC cells. Correspondingly, we observe aberrantly high expression of DARPP-32 proteins in SCLC patient-derived tumor specimens and undetectable levels in normal lung, a finding supported by our genomic analysis of a matched SCLC tumor and adjacent normal tissue RNA-Seq dataset^36^. Genetic ablation of DARPP-32 isoforms in orthotopic SCLC xenografts decreases tumor growth, whereas overexpression of DARPP-32 or t-DARPP in human DMS-53 or H1048 SCLC cells orthotopically injected into mice increases SCLC growth. Collectively, our results suggest upregulation of ASCL1 in SCLC cells transcriptionally activates the DARPP-32 promoter contributing to overexpression of DARPP-32 and t-DARPP proteins, which promote SCLC growth through anti-apoptotic and pro-survival signaling.

Given the primary function of DARPP-32 proteins in neurons and their oncogenic role in non-brain cancers^28^, we hypothesized that DARPP-32 and t-DARPP contribute to neuroendocrine tumor growth in SCLCs. Paul Greengard discovered DARPP-32 as a neuronal phosphoprotein that inhibits PP-1 by mediating dopamine signaling through dopamine D1 and D2 receptors and glutamate signaling through N-methyl-D-aspartate receptor (NMDAR)^19,54–56^. The binding of dopamine, glutamate, nicotine, ethanol, cocaine, and amphetamines to these receptors alters the phosphorylation status of DARPP-32, enabling it to regulate neurotransmission by controlling PP-1 activity^57,58^. Specifically, phosphorylation of DARPP-32 at T34 by PKA inhibits PP-1 activity, whereas Cdk5-mediated phosphorylation of DARPP-32 at T75 abrogates PP-1 action through inhibition of PKA^19^. As a master regulator of PP-1, overexpression of DARPP-32 isoforms in tumor cells enables DARPP-32 and its truncated t-DARPP isoform to control oncogenic cellular functions^8,14,16,17,28,34,59^. For example, t-DARPP, which lacks T34 and the ability to inhibit PP-1, activates pro-survival Akt signaling in breast^20^ and gastric cancers^15^. Likewise, we demonstrated that aberrant upregulation of t-DARPP protein in NSCLC patient-derived tumor tissue correlates with increased lung adenocarcinoma tumor growth (i.e. by T staging score) and that DARPP-32 and t-DARPP promote NSCLC growth via IKKα-induced migration and Akt/Erk-mediated tumor cell survival^8^. Considering our findings in SCLC that DARPP-32 isoforms promote Akt/Erk-mediated proliferation, survival, and protection from apoptosis relative to the collective evidence implicating DARPP-32 and t-DARPP in drug resistance, a future direction beyond the scope of the studies presented here involves investigating the role of DARPP-32 isoforms in SCLC cells acquiring resistance to chemotherapy. Paired chemonaive and chemoresistant SCLC patient-derived xenograft mouse models that would be amenable to addressing this future question have been generated and reported^60^.

The Notch signaling pathway has a complex role in SCLC, but most frequently acts as a tumor suppressor by negatively regulating neuroendocrine differentiation. Activation of Notch signaling is initiated by the interaction of extracellularly expressed canonical delta-like ligands DLL1, DLL3, and DLL4 and jagged ligands (JAG1 and JAG2) with Notch-1, −2, −3, or −4 receptors expressed on the surface of adjacent cells. Ligand binding stimulates receptor cleavage and release of the Notch intercellular domain (NICD), which associates with nuclear transcriptional effectors to regulate transcription of Notch target genes, including activation of hairy and enhancer of split 1 (HES1) and hairy and enhancer of split-related protein 1 (HEY1), transcriptional repressors of ASCL1. The basic helix-loop-helix transcription factor ASCL1 is expressed in approximately 75% of SCLCs, and genetic profiling distinguishes “classic” ASCL1-positive SCLC from “variant” SCLC expressing neurogenic differentiation factor 1 (NEUROD1), another neuronal master regulator that stimulates a neuroendocrine target gene set distinct from that of ASCL1. ASCL1 positively regulates pro-oncogenes associated with SCLC progression and survival, such as Bcl2, SRY-box 2 (SOX2), RET, nuclear factor I B (NFIB). Concordantly with ASCL1 overexpression, Notch is inactivated in most SCLCs by mutations in Notch pathway genes or expression of Notch inhibitors DLK1 and DLL3^30^. A DLL3-targeted antibody-drug conjugate named Rovalpituzumab tesirine (ROVA-T) was shown to have single-agent anti-cancer activity in DLL3-expressing lung neuroendocrine carcinomas and clinical trials in SCLCs are ongoing^61–63^. These results exemplify the highly complex role of Notch signaling in SCLC, which may offer multiple therapeutic targets, and underscore the need to more fully understand the mechanisms of ASLC1-regulated DARPP-32 and Notch signaling in SCLC.

Here, we identify ASCL1 as a transcriptional regulator of DARPP-32. In a subset of SCLC patients with high tumoral t-DARPP expression, we show that the Notch signaling gene set was highly regulated, among which ASCL1 was the top differentially expressed transcript. Expression of ASCL1 and numerous Notch pathway genes was substantially reduced in DARPP-32-depleted human SCLC cells compared to controls. We demonstrate that ASCL1 transcriptionally activates DARPP-32 using reporter assays, site-directed mutagenesis of ASCL1 binding sites on the well-characterized DARPP-32 promoter^42^, and ASCL1 modulation. Interestingly, DARPP-32 may in turn regulate ASCL1 expression through a potential positive feedback mechanism, based on our data showing DARPP-32 knockdown in SCLC cells decreases ASCL1 transcript levels (Fig. 6f). Molecular crosstalk between DARPP-32 isoforms and the Notch signaling pathway in the context of cancer has yet to be reported. However, Notch/RBP-J signaling plays a key role in the regulation of dopamine responsive as neuron-specific loss of Notch/RBP signaling has been shown to cause a deficit in dopamine-dependent instrumental avoidance learning and hyper-responsiveness to apomorphine and SKF38393, a D1 agonist^64^. Brain-derived neurotrophic factor (BDNF) transcriptionally regulates DARPP-32 expression in neurons, as exemplified by delayed and decreased neuronal DARPP-32 expression in BDNF knockout mice, a phenotype that could be rescued *ex vivo* through the addition of BDNF to cultured neurons derived from BDNF null mice^65^. BDNF and ASCL1 were dysregulated and accompanied by aberrant expression of DARPP-32, one of their common transcriptional targets, in striatal neural stem cells derived from a hypoxanthine-guanine phosphoribosyltransferase (HPRT) knockout mouse model of Lesch-Nyhan Syndrome, a neurodevelopmental disorder caused by mutations in the gene encoding HPRT^66^. Based on prior reports in neurons and our evidence in SCLC that ASCL1 regulates DARPP-32 expression, one could surmise that DARPP-32 serves a modulator of neuroendocrine signaling through its control of protein phosphatase activity. Additional studies are necessary to better understand the functional consequences of ASCL1-regulated DARPP-32 activation and to precisely define how DARPP-32 activates downstream signaling, such as Akt, in the context of neuroendocrine tumorigenesis. There is strong collective evidence that t-DARPP activates Akt phosphorylation in cancer, but identification of DARPP-32 and t-DARPP binding partners is necessary to understand exactly how t-DARPP activates Akt through S473 phosphorylation^28^.

SCLC represents a particularly aggressive and deadly form of lung cancer. While the addition of immunotherapy to standard chemotherapy has shown recent promise in the treatment of extensive-stage SCLC^67^, identifying targetable driver mutations and oncogenic pathways in SCLC as the basis for effective molecular targeted therapies has proven challenging^41^. A better molecular understanding SCLC progression will provide a foundation for the discovery of new prognostic tools and therapeutic approaches to improve the clinical outlook and quality of life of patients afflicted with this deadly disease. Here we provide new molecular insights into the mechanisms of SCLC growth. We show that DARPP-32 and t-DARPP promote SCLC proliferation, evasion from caspase-3-dependent apoptosis, activation of Akt and Erk, and increased tumor growth in mice receiving lung xenografts of human SCLC cells stably transduced to overexpress or ablate DARPP-32 isoforms. Patient-derived SCLC tumor specimens exhibit aberrantly high DARPP-32 and t-DARPP protein expression relative to normal lung, in which DARPP-32 isoforms are virtually undetectable by immunostaining. Based on the clinical significance of DARPP-32 and t-DARPP in lung, breast, gastric and other cancers as well as their role in Akt activation and drug resistance, DARPP-32 isoforms may represent a potential therapeutic target^28^. Furthermore, the cancer-specific expression profile of DARPP-32 isoforms lends itself well to safe targeting and minimal toxicity to nonmalignant tissue. In order to develop effective pharmacological inhibitors that target the shared region of DARPP-32 and t-DARPP proteins, the tertiary structure of t-DARPP must be determined. To date, tertiary structures of DARPP-32 isoforms have not been reported because their acidic glutamine- and aspartate-rich central regions likely hinders crystallization of their elongated monomer secondary structure, consisting of 12% alpha helices, 29% beta strands, 24% beta turns, and 35% random coils^68^. However, advances in single-particle cryo-electron microscopy have demonstrated that proteins considerably smaller than the theoretical limit of 50 kDa for cryo-EM can be visualized at near-atomic resolution (i.e. 3.5 to 5 Å), suggesting it may now be technically feasible to solve the tertiary structure of DARPP-32/t-DARPP^69^. Other future directions that will expedite exploration of targeting DARPP-32 and/or t-DARPP signaling as a potential anti-cancer therapy include understanding the exact mechanism(s) through which DARPP-32 isoforms activate Akt and identification of proteins that bind directly to DARPP-32 and/or t-DARPP. It is currently unknown whether most identified upstream and downstream signaling molecules interact with DARPP-32 isoforms directly or indirectly^28^. These future opportunities, the collective prior studies reporting the role of DARPP-32 and t-DARPP in oncogenesis and resistance to therapy, and our findings that upregulation of DARPP-32 isoforms in human SCLC promote growth, anti-apoptotic mechanisms and pro-survival signaling in lung tumor cells of neuroendocrine origin underscore the promise of pursuing DARPP-32 and t-DARPP as potential therapeutic targets or prognostic indicators.

## Methods

### Cell Culture

Human SCLC cell lines, DMS-53 and H1048, were purchased from The European Collection of Authenticated Cell Cultures (ECACC) via Sigma and American Type Culture Collection (ATCC), respectively, and maintained in RPMI-1640 medium (Corning) supplemented with 10% fetal bovine serum (FBS) and antibiotics. Human 293T cells obtained from ATCC were cultured in Dulbecco’s modified Eagle’s medium (DMEM; Corning). All cells were grown in a humidified chamber at 37°C supplied with 5% CO_2_ in media supplemented with 10% fetal bovine serum (FBS; Millipore), 1% Penicillin/Streptomycin antibiotics (Corning), and 25 μg/mL plasmocin (Invivogen). All cell lines were certified prior to purchase and were subsequently authenticated by morphologic inspection on a regular basis.

### Generation of stable cell lines

Expression constructs of human DARPP-32 and t-DARPP cDNA in pcDNA3.1 were generous gifts from Dr. Wael El-Rifai at University of Miami Health System, Florida^17^. The Flag-tagged coding sequences of DARPP-32 and t-DARPP were subcloned into the retroviral vector, pMMP. The pMMP vector and its corresponding pMMP-LacZ control construct were kindly provided by Dr. Debabrata Mukhopadhyay at Mayo Clinic in Jacksonville, Florida^70^. Retrovirus was produced by transfecting 293T cells with packaging plasmids and pMMP vectors encoding DARPP-32 isoforms. The retrovirus was collected at 48h and 72h post-transfection and concentrated using Retro-X concentrator (Takara) according to the manufacturer’s protocol. The concentrated virus was then used to transduce DMS-53 and H1048 cell lines as previously described^71^.

Control pLKO.1-LacZ shRNA and four different lentiviral shRNA pLKO.1 plasmids designed to silence DARPP-32 protein expression were purchased from Sigma. To prepare the lentivirus, shRNA pLKO.1 plasmids along with their corresponding packaging plasmids were transfected in 293T cells. After collecting the media at 48h and 72h post-transfection, the lentivirus was concentrated using Lenti-X concentrator (Takara) according to the manufacturer’s protocol. DMS-53 and H1048 cells transduced with concentrated lentivirus were used for experiments after 72h of puromycin (Sigma) selection.

To determine tumor growth in orthotopic murine models, DMS-53 and H1048 SCLC cells were transduced with lentivirus containing luciferase genes. Briefly, MSCV Luciferase PGK-hygro plasmids were obtained through Dr. Scott Lowe via Addgene (#18782) and transfected in 293T cells with their corresponding packaging plasmids to generate lentivirus. Lenti-X concentrator (Takara) was used to concentrate lentivirus using media at 48h and 72h post-transfection. After transduction with lentivirus following 72h of hygromycin (Sigma) selection, luciferase-labeled stable human SCLC cells were used for experiments.

### Antibodies

Antibodies (200 μg/ml) purchased from Santa Cruz Biotechnology were used to detect DARPP-32 (Polyclonal; Cat no.: sc-11365; Dilution 1:100 and monoclonal; Cat no.: sc-398360; Dilution 1:200) and α-Tubulin (Cat no.: sc-5286; Dilution 1:500). Polyclonal DARPP-32 antibodies were used to generate immunoblotting and immunohistochemistry data in Fig. 1 and Fig. 6, respectively. Alternatively, remaining immunoblotting experiments in Fig. 2 and Fig. 3 were conducted using monoclonal DARPP-32 antibodies. Antibodies (1 μg/μl) against PARP (Cat no.: 9542; Dilution 1:1000), Caspase-3 (Cat no.: 9662; Dilution 1:1000), Cleaved Caspase-3 (Cat no.: 9664; Dilution 1:1000), phosphorylated Akt (S473; Cat no.: 4060; Dilution 1:1000), total Akt (Cat no.: 4691; Dilution 1:1000), phosphorylated p44/42 MAPK (T202/Y204; Cat no.: 4370; Dilution 1:1000) and total p44/42 MAPK (Cat no.: 4695; Dilution 1:1000) were obtained from Cell Signaling Technology. Horseradish peroxidase (HRP)-conjugated anti-rabbit (Cat no.: 7074; Dilution 1:5000) and anti-mouse (Cat no.: 7076; Dilution 1:5000) secondary antibodies (1 μg/μl) purchased from Cell Signaling Technology were used to detect primary antibodies in immunoblotting studies.

### Immunoblotting

DMS-53 and H1048 human SCLC cells lysed in RIPA buffer (Millipore) containing protease inhibitor cocktail (Roche) and phosphatase inhibitor (Millipore) were separated via 4-20% gradient SDS-PAGE (Bio-Rad) and transferred to polyvinyl difluoride membranes (PVDF; Millipore). Membranes blocked with 5% bovine serum albumin (BSA; Sigma) were incubated with primary and secondary antibodies for overnight and 2h, respectively. Antibody-reactive protein bands were detected by adding chemiluminescence substrate (Thermo Fisher Scientific) to the membrane in the dark. The chemiluminescence output was captured electronically using ImageQuant™ LAS 4000 instrument (GE Healthcare).

### Cell growth assay

To measure cell growth by time lapse, 5×10^3^ human SCLC cells were seeded in a 96-well cell culture plate (Cat no.: 655180; Greiner Bio-One). After allowing cells to adhere during a 4h incubation at 37°C, plates were placed in the IncuCyte S3 Live-Cell Analysis System (Essen BioScience). Images of each well were captured with a 4x magnification objective at 8h intervals for up to 72h. Captured images were binarized and further processed using IncuCyte S3 2018B software (Essen BioScience). The segmentation threshold was set to 0.8 to achieve the steepest gradient in intensity between background and foreground cells. Cell confluency (%) was calculated and plotted against time of incubation.

### Cell survival assay

Human SCLC cell lines plated in a 96-well microplate at a concentration of 5×10^3^ cells/well were used to determine cell viability after 72h of incubation using CellTiter 96^®^ AQueous One System (Promega). Epoch microplate spectrophotometer (Biotek) was utilized to record absorbance at 490 nm. The absorbance reading was normalized and plotted as percentage of total viable cells. The average of three independent experiments has been reported.

### Apoptosis analysis

DMS-53 cells transduced with DARPP-32 shRNAs or exogenously overexpressed DARPP-32 isoforms were stained with Annexin V-FITC antibodies (BD Biosciences) to assess apoptosis. To distinguish early apoptotic cells (Annexin-positive and propidium-iodide-negative) from late apoptotic cells (Annexin-positive and propidium-iodide-positive), cells were exposed to propidium iodide (BD Biosciences). Cells undergoing apoptotic cell death were quantified by Annexin V-FITC positivity using flow cytometry.

### Quantitative real-time PCR (qRT-PCR)

Total RNA from DMS-53 cells was extracted using an RNeasy Plus kit (Qiagen) according to the manufacturer's instructions. Isolated RNA (100 ng) was subjected to qRT-PCR analysis using iTaq^™^ Universal SYBR^®^ Green One-Step Kit (Bio-Rad) performed on the 7500 Real-Time PCR System (Applied Biosystems) as previously described^72^. The comparative threshold cycle method (∆∆C_t_) was used to quantify relative amounts of transcripts with *B2M* (β-2-Microglobulin) as an endogenous reference control. Please refer to Supplementary Table 7 for primer sequences.

### BrdU proliferation assay

LacZ control or DARPP-32 shRNA-transduced DMS-53 cells were plated at a density of 1×10^5^ cells per 60-mm plate. After overnight incubation, 30 μM BrdU (Sigma-Aldrich) containing fresh medium was added to the plate for 30 mins. BrdU-labeled cells were then harvested, fixed and incubated with primary monoclonal mouse antibodies (Roche) that recognize bound BrdU in the DNA. BrdU-positive cells were detected using flow cytometry following incubation with secondary FITC-conjugated goat anti-mouse antibody (Invitrogen). Subsequently, propidium iodide staining was performed to differentiate between viable and dead cells. The average of three separate experiments has been documented in the graphs while one representative experiment has been depicted in the main figure.

### Immunohistochemistry

We obtained human small cell lung cancer whole tissue specimens from 6 SCLC patients at Mayo Clinic in Rochester, MN, in accordance with institutional review board-approved protocols. Formalin-fixed. paraffin-embedded whole tissues were serially sectioned, mounted on glass slides, immunostained using a C-terminal antibody that recognize both DARPP-32 and t-DARPP (Santa Cruz Biotechnology, cat no.: sc-11365; 1:100) as previously described^8^. To detect nuclei, slides were counterstained with hematoxylin (IHC World). Tissue microarrays that contained human normal lung tissue from six healthy individuals were purchased from US Biomax Inc. Immunohistochemistry was performed using DARPP-32 antibody as described above. A pulmonary pathologist (A.C.R.) scored each lung tumor specimen by observing the intensity and prevalence of DARPP-32 staining under the microscope.

### RNA-seq data processing

RNA-Seq data of 29 human SCLC patients and 23 corresponding matched normal lung tissue were downloaded from the European Genome-phenome Archive (EGAS00001000334)^36^. All sequencing reads were mapped to the UCSC human reference genome (GRCh38/hg38) using publicly available HISAT2 software with default parameters^73^. Mapped reads were sorted and indexed by SAMtools software^74^. A read summarization function from Subread software, namely featureCounts, was applied to quantify gene expression using transcript per million (TPM)^75^. Using the same methodology, expression of long and short isoforms of *PPP1R1B* gene was calculated. Differential gene expression analysis was performed with R packages, edgeR^76^ and limma^77^. The differentially expressed genes (DEGs) with an absolute value of log_2_FC≥0 and FDR≤0.05 were considered for future analysis. DEGs were ranked based on the highest to lowest fold change values (log_2_FC).

### Pathway Enrichment Analysis

To understand the enriched pathways of the DEGs, Gene Set Enrichment Analysis (GSEA) were performed by GSEA Desktop V3.0 with default parameters using the gene set obtained from c2.cp.kegg.v6.2.symbols.gmt and exported from MSigDB (the Molecular Signatures Database, version 6.2)^78,79^. The RNA-seq read counts of Notch family members enriched in GSEA were identified using same methodology that described above. Heat map generated in GraphPad Prism 8 software was created by plotting log values of RNA-seq read counts for denoted genes involved in Notch signaling.

### In vivo orthotopic lung cancer model

Pathogen-free SCID/NCr mice purchased from the Charles River Laboratories were allowed one week to acclimate to their surroundings, bred, maintained under specific pathogen-free conditions in a temperature-controlled room with alternating 12h light/dark cycles, and fed a standard diet. 1×10^6^ luciferase-labeled human SCLC cells, DMS-53 and H1048, suspended in 80 μl PBS (Corning) and Matrigel^®^ (Corning) were orthotopically injected into the left thorax of 8- to 12-week-old male and female mice. To measure luciferase intensity, mice were imaged after establishment of the lung tumor using an In-Vivo Xtreme xenogen imaging system (Bruker). Bruker molecular imaging software was used to calculate luciferase intensity (Total photons count) of each mouse. Tumor growth was determined by plotting average luciferase intensity over time in GraphPad Prism 8 software. All animal studies were performed in accordance with protocols approved by the University of Minnesota Institutional Animal Care and Use Committee.

### Transient transfection

Human SCLC cell lines, DMS-53 or H1048, were plated in 6-well cell culture plates at a concentration of 1×10^5^ cells per well. The following day, cells were washed with PBS and suspended in OPTI-MEM reduced serum medium (Gibco) for transfection. Polyfect transfection reagent (Qiagen) was used to transfect 1 μg of ASCL1 cDNA plasmids in H1048 cells according to the protocols from the manufacturer. After 4h, antibiotic containing complete RPMI 1640 medium was added to the cells. Cells were harvested, and subsequent experiments were performed 48h post-transfection.

### In vitro promoter luciferase assay

Luciferase reporter of the DARPP-32 promoter was a generous gift from Dr. Wael El-Rifai at University of Miami Health System, Florida^42^. Briefly, investigators amplified the DARPP-32 promoter region (−96 to −3115 from transcription start site) by PCR and then cloned it into the pGL3-basic vector to produce the pGL3- DARPP-32 promoter luciferase reporter. SCLC cells were seeded in 96-well cell culture plates at a density of 1×10^4^ DMS-53 and H1048 cells per well. After 24h post-plating, cells were transfected with 500 ng of pGL3- DARPP-32 plasmids and 50 ng of pRL-SV40 plasmids (renilla; internal reference control) using Polyfect transfection reagent (Qiagen). Lastly, cells were washed with PBS (Corning), lysed with supplied cell culture lysis buffer (Promega), and subjected to dual-luciferase assay after 48h post-transfection using Dual-Luciferase Reporter Assay System (Promega). Luminescence signals from firefly luciferase and renilla luciferase were measured in a multiplate reader, namely Synergy Neo 2 (Biotek).

### Site-directed Mutagenesis

A pair of partially mismatched primers were designed for each ASCL1 binding sites on DARPP-32 promoter using QuikChange Primer Design software (https://www.agilent.com/store/primerDesignProgram.jsp). Primer sequences are available in Supplementary Table 7. Briefly, pGL3-DARPP-32 plasmids were used as templates in PCR reactions to synthesize plasmids containing mutated ASCL1 binding sites by annealing and amplifying site-directed mutagenesis primers using QuikChange Lightning Site-Directed Mutagenesis Kit (Agilent). Digestion with *Dpn* I enzyme was carried out to eliminate methylated parental plasmid strands. Newly synthesized unmethylated plasmids were then transformed into competent *E. coli* XL-10 gold bacteria for nick repair and subsequent amplification. Lastly, plasmids containing mutated ASCL1 binding sites were isolated and verified using Sanger sequencing (Genewiz).

### Statistics

Statistical comparisons performed with one- or two-way analysis of variance (ANOVA) in GraphPad Prism 8 software were considered significant when values of *P* < 0.05. To make comparisons between groups, Dunnett or Sidak multiple comparison tests were performed in all applicable experiments after one- or two-way ANOVA, respectively. Data are expressed as mean ± SEM and representative of at least three independent experiments.

## Data availability

The authors declare that the data supporting the findings of this study are available within the article and its supplementary information.

## Acknowledgements

This study was funded by an NIH/NCI R00-CA187035 grant to L.H.H. as well as support from The Hormel Foundation. We thank Dr. Aaron Mansfield at Mayo Clinic in Rochester, MN for his assistance in the acquisition of human SCLC specimens. We are grateful to Dr. Wael El-Rifai at University of Miami Health System, Florida for generously providing DARPP-32 expression plasmids and DARPP-32 promoter luciferase reporters. We thank Todd Schuster and Josh Monts at The Hormel Institute for their valuable contributions to flow cytometry-based apoptosis and BrdU experiments, Kim Klukas and institutional animal facilities staff for providing excellent animal care and husbandry, and administrative support staff. We appreciate the immunostaining assistance of Mayo Clinic Pathology Research Core.

## Author contributions

S.K.A. conducted in vitro experiments, generated site-directed mutagenesis clones, validated differentially expressed transcripts, performed IncuCyte^®^ live cell imaging, and immunostained normal lung tissue. S.K.A. and L.H.H. managed the mouse colony and performed tumor studies in mice. S.K.A. and L.W. conducted murine in vivo imaging and necropsy. S.K.A. and C.E.H. captured images during immunohistochemistry studies. Y.R. and R.Y. analyzed RNA-seq data, generated differentially expressed gene lists, and performed GSEA using bioinformatics tools. F.K. coordinated acquisition of human SCLC patient tissues and contributed manuscript revisions. A.C.R. pathologically reviewed SCLC patient specimens and scored them based on the staining intensity. S.K.A., L.W., Y.R., R.Y., and L.H.H. contributed to this study by providing critical knowledge to numerous technical aspects of the project. S.K.A. and L.H.H. performed experimental troubleshooting, reviewed relevant scientific literature, critically analyzed data, prepared figures and wrote the manuscript. Conception of aims, project supervision, and acquisition of funding was accomplished by L.H.H. All authors of this manuscript approved the work.

## Competing Financial Interests

The authors declare no competing financial interests.

## Corresponding Author

Correspondence to Luke H. Hoeppner

