## Supplementary Figure 1 for "ASCL1-regulated DARPP-32 and t-DARPP stimulate small cell lung cancer growth and neuroendocrine tumor cell survival"

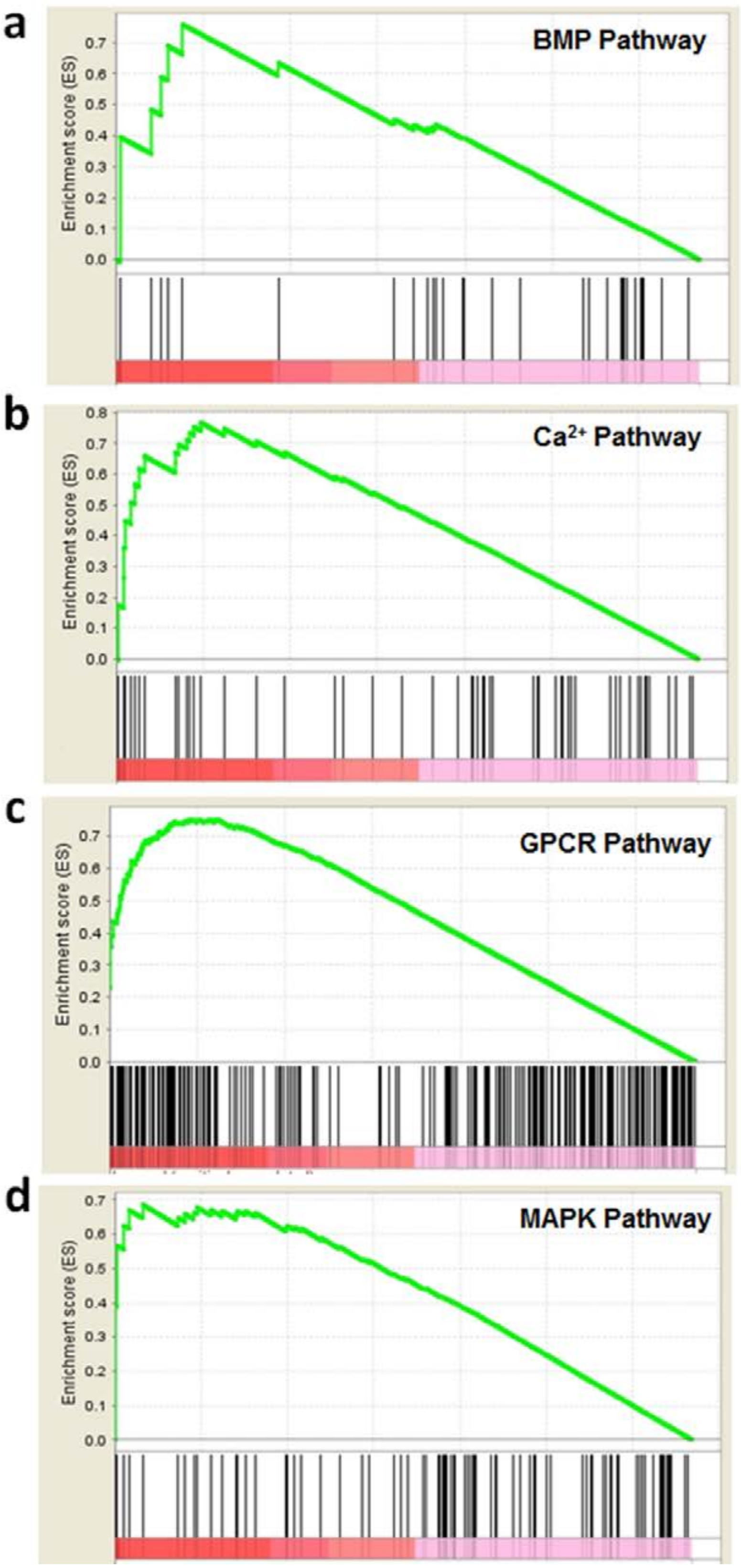


**Supplementary Fig. 1**: Gene set enrichment analysis (GSEA). **a** Bone Morphogenetic Protein (BMP) pathways, **b** Calcium signaling ($Ca^{2+}$) pathways, **c** G protein coupled receptor (GPCR) pathways and **d** mitogen-activated protein kinase (MAPK) pathways were enriched in a subset of SCLC patients with elevated t-DARPP levels in tumor tissue by performing GSEA.
